# Virological characteristics of the SARS-CoV-2 Omicron HK.3 variant harboring the “FLip” substitution

**DOI:** 10.1101/2023.11.14.566985

**Authors:** Yusuke Kosugi, Arnon Plianchaisuk, Olivia Putri, Keiya Uriu, Yu Kaku, Alfredo A. Hinay, Luo Chen, Jin Kuramochi, Kenji Sadamasu, Kazuhisa Yoshimura, Hiroyuki Asakura, Mami Nagashima, The Genotype to Phenotype Japan (G2P-Japan) Consortium, Jumpei Ito, Kei Sato

## Abstract

In November 2023, SARS-CoV-2 XBB descendants, including EG.5.1 (XBB.1.9.2.5.1), the currently predominant lineage, are circulating worldwide according to Nextstrain. EG.5.1 has a characteristic amino acid substitution in the spike protein (S), S:F456L, which contributes to its escape from humoral immunity. EG.5.1 has further evolved, and its descendant lineage harboring S:L455F (i.e., EG.5.1+S:L455F) emerged and was named HK.3 (XBB.1.9.2.5.1.1.3). HK.3 was initially discovered in East Asia and is rapidly spreading worldwide. Notably, the XBB subvariants bearing both S:L455F and S:F456L substitutions, including HK.3, are called the “FLip” variants. These FLip variants, such as JG.3 (XBB.1.9.2.5.1.3.3), JF.1 (XBB.1.16.6.1) and GK.3 (XBB.1.5.70.3), have emerged convergently, suggesting that the acquisition of these two substitutions confers a growth advantage to XBB in the human population. Here, we investigated the virological properties of HK.3 as a representative of the FLip variants.

## Text

In November 2023, SARS-CoV-2 XBB descendants, including EG.5.1 (XBB.1.9.2.5.1), the currently predominant lineage, are circulating worldwide according to Nextstrain (https://nextstrain.org/ncov/gisaid/global/6m). EG.5.1 has a characteristic amino acid substitution in the spike protein (S), S:F456L, which contributes to its escape from humoral immunity (**Figure 1A**).^1^ EG.5.1 has further evolved, and its descendant lineage harboring S:L455F (i.e., EG.5.1+S:L455F) emerged and was named HK.3 (XBB.1.9.2.5.1.1.3) (https://github.com/sars-cov-2-variants/lineage-proposals/issues/414). HK.3 was initially discovered in East Asia and is rapidly spreading worldwide. Notably, the XBB subvariants bearing both S:L455F and S:F456L substitutions, including HK.3, are called the “FLip” variants (https://github.com/cov-lineages/pango-designation). These FLip variants, such as JG.3 (XBB.1.9.2.5.1.3.3), JF.1 (XBB.1.16.6.1) and GK.3 (XBB.1.5.70.3), have emerged convergently, suggesting that the acquisition of these two substitutions confers a growth advantage to XBB in the human population.^2,3^ Here, we investigated the virological properties of HK.3 as a representative of the FLip variants.

**Figure 1.**
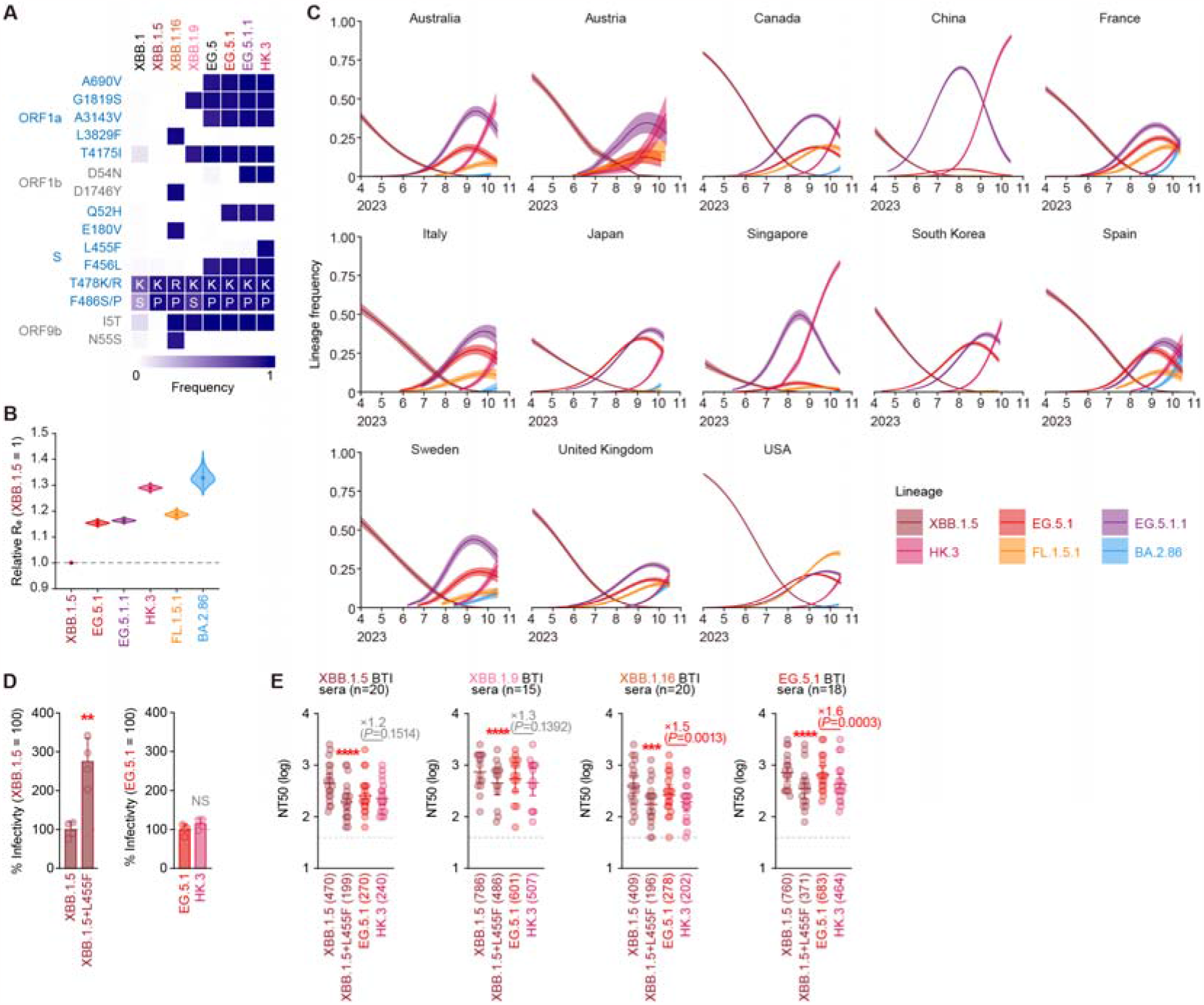
Virological features of HK.3. (A) Frequency of mutations of interest in HK.3 and other lineages of interest. Only mutations with a frequency >0.5 in at least one but not all the representative lineages are shown. **(B)** Estimated global average of relative R_e_ of HK.3 and other lineages of interest. The relative R_e_ of XBB.1.5 is set to 1 (horizontal dashed line). Violin, posterior distribution; dot, posterior mean; line, 95% Bayesian confidence interval. **(C)** Estimated epidemic dynamics of HK.3 and other lineages of interest in the 13 selected countries detected from April 1, 2023 to October 15, 2023. Line, posterior mean; ribbon, 95% Bayesian confidence interval. **(D)** Lentivirus-based pseudovirus assay. HOS-ACE2/TMPRSS2 cells were infected with pseudoviruses bearing each S protein of XBB sublineages. The amount of input virus was normalized to the amount of HIV-1 p24 capsid protein. The percentage infectivity of XBB.1.5, XBB.1.5+L455F compared to that of XBB.1.5 (left), and that of EG.5.1 and HK.3 (EG.5.1+L455F) compared to that of EG.5.1 (right) are shown. The horizontal dash line indicates the mean value of the percentage infectivity of the XBB.1.5 (left) or EG.5.1 (right), respectively. Assays were performed in quadruplicate, and a representative result of six independent assays is shown. The presented data are expressed as the average ± SD. Each dot indicates the result of an individual replicate. Statistically significant differences (**, p < 0.01) versus XBB.1.5 (left) or EG.5.1 (right) were determined by two-sided Student’s *t* tests. Red asterisks indicate increased percentage of infectivity. NS, no statistical significance. **(E)** Neutralization assay. Assays were performed with pseudoviruses harboring the S proteins of XBB.1.5, XBB.1.5+L455F, EG.5.1 and HK.3 (EG.5.1+L455F). The following sera were used: convalescent sera from fully vaccinated individuals who had been infected with XBB.1.5 (eight 3-dose vaccinated donors, seven 4-dose vaccinated donors, four 5-dose vaccinated donors and one 6-dose vaccinated donor. 20 donors in total), XBB.1.9 (three 3-dose vaccinated donors, eight 4-dose vaccinated donors, three 5-dose vaccinated donors and one 6-dose vaccinated donor. 15 donors in total), XBB.1.16 (two 2-dose vaccinated donors, eight 3-dose vaccinated donors, six 4-dose vaccinated donors, two 5-dose vaccinated donors and two 6-dose vaccinated donors. 20 donors in total) and EG.5.1 (one 2-dose vaccinated donor, four 3-dose vaccinated donors, five 4-dose vaccinated donors, four 5-dose vaccinated donors and four 6-dose vaccinated donors. 18 donors in total). Assays for each serum sample were performed in triplicate to determine the 50% neutralization titer (NT_50_). Each dot represents one NT_50_ value, and the geometric mean and 95% confidence interval are shown. The number in parenthesis indicates the geometric mean of NT_50_ values. The horizontal dash line indicates the detection limit (40-fold). Statistically significant differences versus XBB.1.5 (***, p < 0.001, ****, p < 0.0001) or versus EG.5.1 were determined by two-sided Wilcoxon signed-rank tests. Red asterisks indicate decreased NT_50_s. The fold change of reciprocal NT_50_ is calculated between EG.5.1 and HK.3. Background information on the convalescent donors is summarized in **Table S1**.

We estimated the relative effective reproduction number (R_e_) of HK.3 based on the genome surveillance data from thirteen countries with significant presence of HK.3 using a Bayesian hierarchical multinomial logistic regression model (**Figure 1B, Tables S2 and S3**).^4^ The global average of R_e_ of HK.3 is 1.29- and 1.12-fold higher than that of XBB.1.5 and EG.5.1, suggesting that HK.3 potentially become the predominant lineage worldwide soon. Indeed, as of the beginning of October 2023, HK.3 have already outcompeted EG.5.1 in some countries such as Australia, China, South Korea, and Singapore (**Figure 1C**).

Next, to assess the possibility that the enhanced infectivity of HK.3 contributes to its augmented R_e_, we prepared the lentivirus-based pseudoviruses with the S proteins of XBB.1.5, EG.5.1, HK.3 and an XBB.1.5 derivative, XBB.1.5+L455F. Although the S:L455F substitution significantly increased infectivity of XBB.1.5, the infectivity of HK.3 (identical to EG.5.1+S:L455F) was comparable to that of EG.5.1 (**Figure 1D**). The discrepancy of the effect of S:L455F between XBB.1.5 and EG.5.1 might be attributed to the epistatic effect of structures of the S proteins of XBB.1.5 and EG.5.1. These results suggest that the increased R_e_ of HK.3 is not due to the increased infectivity caused by S:L455F.

We then performed a neutralization assay using the breakthrough infection (BTI) sera infected with XBB.1.5, XBB.1.9, XBB.1.16 or EG.5.1 to address whether HK.3 evades the antiviral effect of humoral immunity induced by BTI of these variants. The 50% neutralization titer (NT_50_) of all BTI sera tested against XBB.1.5+S:L455F was significantly lower than those against the parental XBB.1.5 (**Figure 1E**). Notably, the NT_50_ of EG.5.1 BTI sera against HK.3 was significantly (1.6-fold, p=0.0003) lower than that against EG.5.1 (**Figure 1E**). These results suggest that the increased R_e_ of HK.3 is partly attributed to the immune evasion from the humoral immunity elicited by the BTI of XBB subvariants including EG.5.1, its ancestor, and S:L455F is a key mutation leading to this immune evasion.

## Grants

Supported in part by AMED SCARDA Japan Initiative for World-leading Vaccine Research and Development Centers “UTOPIA” (JP223fa627001, to Kei Sato), AMED SCARDA Program on R&D of new generation vaccine including new modality application (JP223fa727002, to Kei Sato); AMED Research Program on Emerging and Re-emerging Infectious Diseases (JP22fk0108146, to Kei Sato; JP21fk0108494 to G2P-Japan Consortium and Kei Sato; JP21fk0108425, to Kei Sato; JP21fk0108432, to Kei Sato; JP22fk0108511, to G2P-Japan Consortium and Kei Sato; JP22fk0108516, to Kei Sato; JP22fk0108506, to Kei Sato); AMED Research Program on HIV/AIDS (JP22fk0410039, to Kei Sato); JST PRESTO (JPMJPR22R1, to Jumpei Ito); JST CREST (JPMJCR20H4, to Kei Sato); JSPS KAKENHI Grant-in-Aid for Early-Career Scientists (23K14526, to Jumpei Ito); JSPS Core-to-Core Program (A. Advanced Research Networks) (JPJSCCA20190008, Kei Sato); JSPS Research Fellow DC2 (22J11578, to Keiya Uriu); JSPS Research Fellow DC1 (23KJ0710, to Yusuke Kosugi); The Tokyo Biochemical Research Foundation (to Kei Sato); The Mitsubishi Foundation (to Kei Sato).

## Declaration of interest

J.I. has consulting fees and honoraria for lectures from Takeda Pharmaceutical Co. Ltd. K.S. has consulting fees from Moderna Japan Co., Ltd. and Takeda Pharmaceutical Co. Ltd. and honoraria for lectures from Gilead Sciences, Inc., Moderna Japan Co., Ltd., and Shionogi & Co., Ltd. The other authors declare no competing interests. All authors have submitted the ICMJE Form for Disclosure of Potential Conflicts of Interest. Conflicts that the editors consider relevant to the content of the manuscript have been disclosed.

