## Supplementary Appendix for "Virological characteristics of the SARS-CoV-2 Omicron HK.3 variant harboring the “FLip” substitution"

**Table of Contents**

**Contents** Page

**Materials and Methods** 2-5
Ethics statement
Human serum collection
Mutation frequency calculation and epidemic dynamics analysis
Plasmid construction
Cell culture
Pseudovirus preparation
Neutralization assay
Data availability
**Table S1.** Human sera used in this study 6-8
**Table S2.** Estimated global average of relative $R_e$ and related epidemic dynamics modeling parameters of SARS-CoV-2 lineages of interest circulated in 13 countries from April 1, 2023 to October 15, 2023 9
**Table S3.** Estimated national relative $R_e$ and related epidemic dynamics modeling parameters of SARS-CoV-2 lineages of interest circulated in 13 countries from April 1, 2023 to October 15, 2023 10-14
**Table S4.** Primers used in this study 15
**Consortia** 16
**Acknowledgments** 17
**Supplemental References** 18

### Materials and Methods

#### Ethics statement

All protocols involving specimens from human subjects recruited at Interpark Kuramochi Clinic was reviewed and approved by the Institutional Review Board of Interpark Kuramochi Clinic (approval ID: G2021-004). All human subjects provided written informed consent. All protocols for the use of human specimens were reviewed and approved by the Institutional Review Boards of The Institute of Medical Science, The University of Tokyo (approval IDs: 2021-1-0416 and 2021-18-0617).

#### Human serum collection

Convalescent sera were collected from fully vaccinated individuals who had been infected with XBB.1.5 (eight 3-dose vaccinated, seven 4-dose vaccinated, four 5-dose vaccinated and one 6-dose vaccinated; time interval between the last vaccination and infection, 44-524 days; 14-46 days after testing. n=20 in total; average age: 46.3 years, range: 15-74 years, 30% male), XBB.1.9 (three 3-dose vaccinated, eight 4-dose vaccinated, three 5-dose vaccinated and one 6-dose vaccinated; time interval between the last vaccination and infection, 19–535 days; 6–32 days after testing. n=15 in total; average age: 55 years, range: 32–88 years, 53.3% male), XBB.1.16 (two 2-dose vaccinated, eight 3-dose vaccinated, six 4-dose vaccinated, two 5-dose vaccinated and two 6-dose vaccinated; time interval between the last vaccination and infection, 20–653 days; 7–30 days after testing. n=20 in total; average age: 48.3 years, range: 18–86 years, 50% male) and EG.5.1 (one 2-dose vaccinated, four 3-dose vaccinated, five 4-dose vaccinated, four 5-dose vaccinated and four 6-dose vaccinated; time interval between the last vaccination and infection, 58–500 days; 4–27 days after testing. n=18 in total; average age: 55.1 years, range: 27–77 years, 50% male). The SARS-CoV-2 variants were identified as previously described.[1-3] Sera were inactivated at 56°C for 30 minutes and stored at –80°C until use. The details of the convalescent sera are summarized in **Table S1**.

#### Mutation frequency calculation and epidemic dynamics analysis

We modeled the epidemic dynamics of SARS-CoV-2 with the GISAID viral genomic surveillance data (https://www.gisaid.org/; downloaded on October 27, 2023)[4]. The PANGO lineage of each SARS-CoV-2 isolate was reassigned using NextClade v2.14.0[5]. We discarded the data of SARS-CoV-2 that i) was isolated from non-human hosts; ii) was sampled from the original passage; iii) lacks NextClade PANGO lineage information; and iv) whose genomic sequence is not longer than 28,000 base pairs and contains ≥2% of unknown (N) nucleotides.

We randomly selected at most 500 genomic sequences of SARS-CoV-2 in XBB.1, XBB.1.5, XBB.1.16, XBB.1.9, EG.5, EG.5.1, EG.5.1.1, and HK.3 for calculating a mutation frequency (EPI SET ID: EPI_SET_231111fo). A mutation frequency of each subvariant Is calculated by dividing the number of sequences harboring the mutation of interest with the total number of sequences in that subvariant.

To estimate the national $R_e$ and the global average for HK.3 and other lineages of interest, we analyzed data of SARS-CoV-2 isolated circulating from April 1, 2023 to October 15, 2023. We selected 13 countries with 5,000 SARS-CoV-2 isolates in total and picked only lineages having ≥20 isolates in all selected countries, resulting in data of 145,615 SARS-CoV-2 isolates in total (EPI SET ID: EPI_SET_231110rt). The countries include Australia, Austria, Canada, China, France, Italy, Japan, Singapore, South Korea, Spain, Sweden, the UK, and the USA. The selected lineages include EG.1, EG.5.1, EG.5.1.1, FL.1.5.1, FL.2, FL.4, FL.5, FU.1, HK.3, XBB.1.16. XBB.1.16.1, XBB.1.5, XBB.1.5.24, XBB.1.9.1, XBB.1.9.2, and XBB.2.3. We pooled BA.2.86 and BA.2.86.1 together as BA.2.86 and additionally included BA.2.86 in the list of selected lineages. Then, we calculated the daily count of SARS-CoV-2 isolates in each lineage of each country and employed a hierarchical Bayesian multinomial logistic model, established in our previous study[6], to estimate the national $R_e$ and the global average for each lineage. Briefly, the national relative $R_e$ of each lineage $l$ of a country $c$ ($r_{lc}$) was calculated according to the national slope parameter $\beta_{lc}$ as $r_{lc} = \exp(\gamma\beta_{lc})$ where $\gamma$ is the average viral generation time (2.1 days) (http://sonorouschocolate.com/covid19/index.php?title=Estimating_Generation_Time_Of_Omicron). Similarly, the global average of relative $R_e$ of each lineage was calculated according to the global slope parameter $\beta_l$ as $r_l = \exp(\gamma\beta_l)$. The global intercept and slope parameters of XBB.1.5 were fixed at 0. The relative $R_e$ of XBB.1.5 was fixed at 1, and that of the other lineages was estimated relatively to that of XBB.1.5. Parameter estimation was performed using the Markov Chain Monte Carlo (MCMC) approach implemented in CmdStan v2.33.1 provided from CmdStanr R package v0.6.1 (https://mc-stan.org/cmdstanr/). Four independent MCMC chains were run for 5,000 iterations including 1,000 warmup iterations. We confirmed that an estimated $\hat{R}$ diagnostic value is <1.01 and bulk and tail effective sampling size are >200 for all runs, indicating that they were successfully convergent. Information on the estimated parameters is summarized in **Tables S2 and S3**. Only the results for XBB.1.5, EG.5.1, EG.5.1.1, HK.3, FL.1.5.1, and BA.2.86 are shown in **Figures S1B and S1C**.

**Plasmid construction**

Plasmids expressing the SARS-CoV-2 spike proteins of Omicron XBB.1.5, EG.5.1 and its derivative were prepared in our previous studies.[7,8] Plasmids expressing the spike protein of HK.3 and its derivative were generated by site-directed overlap extension PCR using pC-SARS2-S XBB.1.5[7] or pC-SARS2-S EG.5.1[8] as the template and the primers listed in **Table S4**. The resulting PCR fragment was subcloned into the KpnI-NotI site of the pCAGGS vector[9] using In-Fusion HD Cloning Kit (Takara, Cat# Z9650N). Nucleotide sequences were determined by DNA sequencing services (Eurofins), and the sequence data were analyzed by SnapGene software v6.1.1 (www.snapgene.com).

**Cell culture**

The Lenti-X 293T cell line (Takara, Cat# 632180) and HOS-ACE2/TMPRSS2 cells (kindly provided by Dr. Kenzo Tokunaga), a derivative of HOS cells (a human osteosarcoma cell line; ATCC CRL-1543) stably expressing human ACE2 and TMPRSS2,[10,11] were maintained in Dulbecco's modified Eagle's medium (DMEM) (high glucose) (Wako, Cat# 044- 29765) containing 10% fetal bovine serum (Sigma-Aldrich Cat# 172012-500ML), 100 units penicillin and 100 ug/ml streptomycin (Sigma-Aldrich, Cat# P4333-100ML).

**Pseudovirus preparation**

Pseudoviruses were prepared as previously described.[12] Briefly, lentivirus (HIV-1)-based, luciferase-expressing reporter viruses were pseudotyped with the SARS-CoV-2 spikes (S). One prior day of transfection, the LentiX-293T cells were seeded at a density of 2 × $10^6$ cells. The LentiX-293T cells were cotransfected with 1 μg psPAX2-IN/HiBiT (a packaging plasmid encoding the HiBiT-tag-fused integrase[13]), 1 μg pWPI-Luc2 (a reporter plasmid encoding a firefly luciferase gene[13]) and 500 ng plasmids expressing parental S or its derivatives using TransIT-293 transfection reagent (Mirus, Cat# MIR2704) according to the manufacturer's protocol. Two days post transfection, the culture supernatants were harvested and filtrated. The amount of produced pseudovirus particles was quantified by the HiBiT assay using Nano Glo HiBiT lytic detection system (Promega, Cat# N3040) as previously described[13]. In this system, HiBiT peptide is produced with HIV-1 integrase and forms NanoLuc luciferase with LgBiT, which is supplemented with substrates. In each pseudovirus particle, the detected HiBiT value is correlated with the amount of the pseudovirus capsid protein, HIV-1 p24 protein.[13] Therefore, we calculated the amount of HIV-1 p24 capsid protein based on the HiBiT value measured, according to the previous paper.[13] To measure viral infectivity, the same amount of pseudovirus normalized with the HIV-1 p24 capsid protein was inoculated into HOS-ACE2/TMPRSS2 cells. At two days postinfection, the infected cells were lysed with a Bright-Glo luciferase assay system (Promega, Cat# E2620), and the luminescent signal produced by firefly luciferase reaction was measured using a GloMax explorer multimode microplate reader 3500 (Promega). The pseudoviruses were stored at –80°C until use.

**Neutralization assay**

Neutralization assays were performed as previously described.[12] The SARS-CoV-2 spike pseudoviruses (counting ~100,000 relative light units) were incubated with serially diluted (40-fold to 29,160-fold dilution at the final concentration) heat-inactivated sera at 37°C for 1 hour. Pseudoviruses without sera were included as controls. Then, 20 μl mixture of pseudovirus and serum was added to HOS-ACE2/TMPRSS2 cells (10,000 cells/100 μl) in a 96-well white plate. Two days post infection, the infected cells were lysed with a Bright-Glo luciferase assay system (Promega, Cat# E2620), and the luminescent signal was measured using a GloMax explorer multimode microplate reader 3500 (Promega). The assay of each serum sample was performed in triplicate, and the 50% neutralization titer ($NT_{50}$) was calculated using Prism 9 (GraphPad Software).

**Data availability**

Dataset used in the epidemic dynamics analysis in this study is available from the GISAID database (https://www.gisaid.org; EPI_SET_231110rt and EPI_SET_231111fo). The GISAID supplemental tables for EPI_SET_230725pv is available in the GitHub repository （https://github.com/TheSatoLab/HK.3_short).

**Table S1. Human sera used in this study**

| SARS-CoV-2 infected | Donor ID | Sex | Age | Date of 1st vaccination (YYYY-MM-DD) | Date of 2nd vaccination (YYYY-MM-DD) | Date of 3rd vaccination (YYYY-MM-DD) | Date of 4th vaccination (YYYY-MM-DD) | Date of 5th vaccination (YYYY-MM-DD) | Date of 6th vaccination (YYYY-MM-DD) | Date of test (YYYY-MM-DD) | Date of sampling (YYYY-MM-DD) | Prior infection? |
|---|---|---|---|---|---|---|---|---|---|---|---|---|
| XBB.1.5 | 37306 | Female | 53 | 2021-04-27 (P) | 2021-05-18 (P) | 2022-02-01 (P) | 2022-07-30 (M) | 2022-12-17 (P) | | 2023-07-20 | 2023-08-08 | No |
| XBB.1.5 | 37097 | Female | 44 | NA (M) | 2021-08-16 (M) | 2022-05-13 (M) | | | | 2023-07-13 | 2023-08-11 | No |
| XBB.1.5 | 37598 | Female | 43 | 2021-04-28 (P) | 2021-05-19 (P) | 2022-11-08 (P) | 2022-07-08 (M) | 2022-12-27 (P) | | 2023-07-28 | 2023-08-11 | Yes |
| XBB.1.5 | 37072 | Female | 15 | 2021-09-25 (P) | 2021-10-18 (P) | 2022-05-02 (P) | | | | 2023-07-11 | 2023-08-11 | No |
| XBB.1.5 | 37071 | Female | 48 | 2021-10-01 (P) | 2021-11-01 (P) | 2022-05-06 (P) | | | | 2023-07-15 | 2023-08-11 | No |
| XBB.1.5 | 36845 | Male | 29 | 2021-09-01 (M) | 2021-09-29 (M) | 2022-05-27 (M) | | | | 2023-06-26 | 2023-08-11 | No |
| XBB.1.5 | 37229 | Female | 74 | 2021-06-24 (P) | 2021-07-15 (P) | 2022-02-16 (M) | 2022-07-20 (M) | 2023-03-25 (M) | | 2023-07-17 | 2023-08-01 | No |
| XBB.1.5 | 36708 | Male | 55 | 2021-08-07 (P) | 2021-08-27 (P) | 2022-04-14 (P) | | | | 2023-06-01 | 2023-07-09 | No |
| XBB.1.5 | 38084 | Male | 44 | 2021-09-13 (P) | 2021-10-05 (P) | 2022-07-29 (M) | | | | 2023-08-11 | 2023-09-02 | No |
| XBB.1.5 | 37998 | Female | 65 | 2021-08-04 (P) | 2021-08-30 (P) | 2022-03-19 (M) | 2022-09-02 (P) | 2022-12-24(PBA.4/5) | 2023-06-27 (P) | 2023-08-10 | 2023-09-02 | No |
| XBB.1.5 | 37798 | Female | 62 | 2021-03-17 (P) | 2021-04-09 (P) | 2021-12-23 (P) | 2022-07-28 (P) | 2023-06-17 (P) | | 2023-08-03 | 2023-08-20 | No |
| XBB.1.5 | 38061 | Female | 55 | 2021-08-17 (P) | 2021-09-18 (P) | 2022-04-02 (M) | 2022-10-14(PBA.1) | | | 2023-08-11 | 2023-09-04 | No |
| XBB.1.5 | 38019 | Female | 18 | 2021-09-07 (P) | 2021-10-07 (P) | 2022-04-28 (P) | 2022-12-27 (P) | | | 2023-08-10 | 2023-09-04 | No |
| XBB.1.5 | 38952 | Female | 54 | 2021-07-27 (M) | 2021-08-24 (M) | 2022-03-24 (P) | 2022-10-26 (P) | | | 2023-08-23 | 2023-09-10 | No |
| XBB.1.5 | 38871 | Male | 46 | 2021-09-12 (P) | 2021-10-03 (P) | 2022-04-08 (M) | 2022-11-05 (M) | | | 2023-08-22 | 2023-09-10 | No |
| XBB.1.5 | 38880 | Male | 51 | 2021-10-07 (P) | 2021-10-28 (P) | 2022-05-13 (M) | 2022-11-09 (PBA.4/5) | | | 2023-08-22 | 2023-09-16 | No |
| XBB.1.5 | 39018 | Female | 51 | 2021-07-29 (P) | 2021-08-23 (P) | 2022-03-19 (P) | | | | 2023-08-25 | 2023-09-13 | Yes |
| XBB.1.5 | 39019 | Female | 22 | 2021-07-26 (M) | 2021-08-23 (M) | 2022-03-19 (P) | | | | 2023-08-25 | 2023-09-13 | No |
| XBB.1.5 | 39293 | Female | 49 | 2021-05-11 (P) | 2021-06-01 (P) | 2022-02-15 (M) | 2022-09-16 (M) | | | 2023-08-31 | 2023-09-21 | No |
| XBB.1.5 | 39296 | Male | 48 | 2021-07-15 (M) | 2021-08-23 (M) | 2022-03-24 (M) | 2022-12-25 (M) | | | 2023-09-01 | 2023-09-23 | No |
| XBB.1.9 | 37317 | Male | 65 | NA (P) | NA (P) | NA (P) | NA (P) | NA (P) | | 2023-07-20 | 2023-08-12 | No |
| XBB.1.9 | 37087 | Female | 50 | 2021-09-05 (P) | 2021-09-26 (P) | 2022-03-26 (M) | 2022-11-05 (P) | | | 2023-07-11 | 2023-08-12 | No |
| XBB.1.9 | P592 | Female | 32 | NA (NA) | NA (NA) | 2022-10-26 (P) | | | | 2023-05-06 | 2023-05-16 | No |
| XBB.1.9 | P595 | Female | 88 | 2021-05-18 (P) | 2021-06-08 (P) | 2022-02-04 (M) | 2022-07-19 (M) | 2022-11-18 (P) | | 2023-05-13 | 2023-05-19 | No |
| XBB.1.9 | KS-230718 | Male | 41 | 2021-06-17 (P) | 2021-07-07 (P) | 2022-03-28 (M) | 2022-10-27(MBA.4/5) | | | 2023-06-30 | 2023-07-18 | No |
| XBB.1.9 | 38752 | Male | 54 | 2021-10-02 (P) | 2021-11-02 (P) | 2022-05-27 (M) | 2022-12-02(Novavax) | | | 2023-08-20 | 2023/9/13 | No |
| XBB.1.9 | 38046 | Female | 59 | 2021-08-25 (P) | 2021-09-18 (P) | 2022-04-01 (M) | 2022-10-12 (P) | | | 2023-08-11 | 2023-08-30 | No |
| XBB.1.9 | 38365 | Female | 54 | 2021-07-13 (P) | 2021-08-04 (P) | 2022-02-26 (P) | | | | 2023-08-15 | 2023-08-29 | No |

| | | | | | | | | | | | | |
|---|---|---|---|---|---|---|---|---|---|---|---|---|
| XBB.1.9 | 37744 | Female | 43 | 2021-09-07 (P) | 2021-10-05 (P) | 2022-04-12 (M) | 2022-10-24 (P) | | | 2023-08-01 | 2023-08-29 | No |
| XBB.1.9 | 38926 | Male | 49 | 2021-06-01 (P) | 2021-06-22 (P) | 2022-03-21 (M) | | | | 2023-08-23 | 2023-09-11 | No |
| XBB.1.9 | 39015 | Male | 54 | 2021-08-08 (P) | 2021-08-29 (P) | 2022-03-19 (M) | 2022-12-17 (P) | | | 2023-08-25 | 2023-09-16 | No |
| XBB.1.9 | 39035 | Male | 64 | 2021-07-14 (P) | 2021-08-05 (P) | 2022-03-22 (P) | 2022-09-08 (M) | 2022-12-19 (P) | 2023-08-07 (P) | 2023-08-26 | 2023-09-14 | No |
| XBB.1.9 | 39317 | Male | 59 | 2021-09-03 (P) | 2021-09-24 (P) | 2022-03-27 (P) | 2022-10-22 (P) | | | 2023-09-01 | 2023-09-23 | No |
| XBB.1.9 | 39338 | Male | 55 | 2021-08-26 (P) | 2021-10-07 (P) | 2022-04-09 (M) | 2022-10-15 (P) | | | 2023-08-31 | 2023-09-23 | No |
| XBB.1.9 | 39446 | Female | 58 | 2021-04-27 (P) | 2021-05-18 (P) | 2022-01-21 (P) | 2022-12-14 (MBA.4/5) | 2023-06-03 (M) | | 2023-09-04 | 2023-09-27 | No |
| XBB.1.16 | KY-230807 | Female | 53 | 2021-08-18 (P) | 2021-09-08 (P) | 2022-04-13 (P) | 2022-10-21 (P) | | | 2023-07-24 | 2023-08-07 | No |
| XBB.1.16 | LC-230807 | Male | 26 | 2021-07-22 (P) | 2021-08-21 (P) | 2022-03-30 (M) | | | | 2023-07-24 | 2023-08-07 | No |
| XBB.1.16 | 37090 | Male | 56 | 2021-08-01 (P) | 2021-08-22 (P) | 2022-04-08 (P) | 2022-09-24 (P) | 2023-01-20 (P) | 2023-06-07 (P) | 2023-07-12 | 2023-08-11 | No |
| XBB.1.16 | 37300 | Female | 54 | 2021-08-31 (P) | 2021-09-29 (P) | 2022-04-08 (P) | 2022-09-28 (P) | | | 2023-07-19 | 2023-08-12 | No |
| XBB.1.16 | 37302 | Female | 42 | 2021-09-06 (M) | 2021-10-04 (M) | | | | | 2023-07-19 | 2023-08-12 | No |
| XBB.1.16 | AH-230816 | Male | 34 | 2021-08-27 (M) | 2021-09-24 (M) | 2022-04-16 (M) | | | | 2023-08-03 | 2023-08-16 | Yes |
| XBB.1.16 | P601 | Male | 38 | 2021-11-05 (P) | 2021-11-26 (P) | 2022-05-27 (P) | | | | 2023-07-06 | 2023-07-14 | No |
| XBB.1.16 | P593 | Male | 65 | 2021-08-02 (P) | 2021-08-30 (P) | 2022-04-03 (P) | | | | 2023-05-09 | 2023-05-16 | No |
| XBB.1.16 | P594 | Male | 18 | 2021-09-06 (P) | 2021-10-15 (P) | | | | | 2023-05-07 | 2023-05-19 | No |
| XBB.1.16 | JI-230718 | Male | 34 | 2021-07-19 (M) | 2021-08-20 (M) | 2022-05-13 (M) | 2022-11-22 (M) | | | 2023-07-03 | 2023-07-18 | Yes |
| XBB.1.16 | 38047 | Female | 37 | 2021-08-23 (M) | 2021-09-20 (M) | 2022-03-28 (M) | 2022-09-22 (M) | 2023-02-09 (P) | | 2023-08-11 | 2023-08-30 | No |
| XBB.1.16 | 38078 | Male | 32 | 2021-09-08 (M) | 2021-10-06 (M) | 2022-04-15 (M) | 2022-10-14 (M) | | | 2023-08-11 | 2023-09-02 | No |
| XBB.1.16 | 38397 | Female | 55 | 2021-06-03 (P) | 2021-06-24 (P) | 2022-01-18 (P) | 2022-06-28 (P) | | | 2023-08-15 | 2023-09-02 | No |
| XBB.1.16 | 38060 | Male | 46 | 2021-09-06 (P) | 2021-09-27 (P) | 2022-05-17 (P) | | | | 2023-08-11 | 2023-09-02 | No |
| XBB.1.16 | 37324 | Male | 57 | NA (P) | NA (P) | NA (P) | | | | 2023-07-20 | 2023-08-16 | No |
| XBB.1.16 | 38016 | Female | 65 | 2021-07-13 (P) | 2021-08-03 (P) | 2022-03-14 (M) | 2022-08-19 (M) | 2022-12-19 (PBA.4/5) | 2023-07-21 (PBA.4/5) | 2023-08-10 | 2023-08-29 | No |
| XBB.1.16 | 39010 | Female | 86 | 2021-06-01 (P) | 2021-06-22 (P) | 2022-01-29 (M) | 2022-07-25 (NA) | 2023-11-13 (NA) | | 2023-08-25 | 2023-09-08 | No |
| XBB.1.16 | 39030 | Female | 65 | 2021-07-28 (P) | 2021-09-01 (P) | 2022-03-15 (M) | 2022-12-16 (P) | | | 2023-08-26 | 2023-09-10 | No |
| XBB.1.16 | 38090 | Female | 62 | 2021-08-28 (NA) | 2021-09-18 (NA) | 2022-07-23 (NA) | | | | 2023-08-11 | 2023-09-05 | No |
| XBB.1.16 | P610 | Female | 40 | 2021-08-05 (P) | 2021-08-26 (P) | 2022-03-29 (M) | | | | 2023-08-08 | 2023-09-07 | No |
| EG.5 | P618 | Male | 54 | 2021-08-05 (P) | 2021-08-26 (P) | 2022-03-17 (P) | 2022-08-24 (P) | 2022-11-24 (P) | | 2023-08-23 | 2023-09-07 | No |
| EG.5 | KK-230801 | Male | 56 | 2021-07-16 (P) | 2021-08-06 (P) | 2022-03-03 (M) | 2022-08-09 (P) | | | 2023-07-28 | 2023-08-01 | No |
| EG.5 | 37301 | Female | 27 | 2021-08-01 (M) | 2022-03-06 (M) | | | | | 2023-07-19 | 2023-08-12 | No |
| EG.5 | 37330 | Female | 42 | 2021-09-25 (P) | 2021-10-16 (P) | 2022-05-13 (P) | | | | 2023-07-20 | 2023-08-13 | No |
| EG.5 | 38111 | Female | 50 | 2021-08-16 (P) | 2021-09-06 (P) | 2022-04-01 (P) | | | | 2023-08-12 | 2023-09-02 | No |

| | | | | | | | | | | | | |
|---|---|---|---|---|---|---|---|---|---|---|---|---|
| EG.5 | 38197 | Female | 49 | 2021-08-20 (P) | 2021-09-10 (P) | 2022-03-25 (M) | 2022-11-18 (P) | | | 2023-08-13 | 2023-09-02 | No |
| EG.5 | 38217 | Male | 62 | 2021-07-04 (P) | 2021-07-25 (P) | 2022-02-26 (M) | 2022-08-06 (M) | 2022-11-20 (P) | | 2023-08-13 | 2023-09-02 | No |
| EG.5 | 37999 | Male | 67 | 2021-06-07 (P) | 2021-07-01 (P) | 2022-02-08 (P) | 2022-07-12 (P) | 2022-12-24 (PBA.4/5) | 2023-06-13 (MBA.4/5) | 2023-08-10 | 2023-09-02 | No |
| EG.5 | 38946 | Female | 65 | 2021-07-17 (P) | 2021-08-07 (P) | 2022-03-06 (M) | 2022-08-06 (M) | 2022-11-25 (P) | | 2023-08-23 | 2023-09-10 | No |
| EG.5 | 39025 | Male | 63 | 2021-08-01 (P) | 2021-08-22 (P) | 2022-03-07 (M) | 2022-08-10 (M) | 2022-11-26 (PBA.4/5) | 2023-06-06 (PBA.4/5) | 2023-08-26 | 2023-09-11 | No |
| EG.5 | 39288 | Male | 77 | 2021-06-09 (P) | 2021-07-01 (P) | 2022-02-05 (M) | 2022-07-12 (P) | 2022-11-18 (PBA.4/5) | 2023-05-30 (PBA.1) | 2023-08-31 | 2023-09-20 | No |
| EG.5 | 39301 | Male | 41 | 2021-08-26 (M) | 2021-09-23 (M) | 2022-04-24 (M) | 2022-10-09 (PBA.1) | | | 2023-09-01 | 2023-09-23 | No |
| EG.5 | 39314 | Female | 56 | 2021-07-25 (P) | 2021-08-22 (P) | 2022-03-03 (M) | 2022-08-06 (M) | 2022-11-06 (M) | 2023-04 (NA) | 2023-09-01 | 2023-09-21 | No |
| EG.5 | 39315 | Female | 60 | 2021-08-29 (P) | 2021-09-19 (P) | 2022-04-16 (M) | 2022-09-16 (P) | 2022-12-16 (PBA.4/5) | | 2023-08-31 | 2023-09-23 | No |
| EG.5 | 39316 | Male | 63 | 2021-07-31 (P) | 2021-08-21 (P) | 2022-04-23 (M) | 2022-09-30 (PBA.1) | | | 2023-08-31 | 2023-09-23 | No |
| EG.5 | 39330 | Male | 59 | 2021-08-24 (P) | 2021-09-14 (P) | 2022-03-26 (P) | 2022-10-23 (M) | | | 2023-08-31 | 2023-09-23 | No |
| EG.5 | 39503 | Female | 41 | 2021-08-24 (P) | 2021-09-15 (P) | 2022-04-27 (P) | | | | 2023-09-05 | 2023-09-25 | No |
| EG.5 | 39328 | Female | 59 | 2021-09-30 (P) | 2021-10-27 (P) | 2022-05-17 (P) | | | | 2023-08-31 | 2023-09-27 | Yes |

NA, not applicable.

P, Pfizer-BioNTech; M, Moderna

**Table S2. Estimated global average of relative $R_e$ and related epidemic dynamics modeling parameters of the SARS-CoV-2 lineages of interest circulated in 13 countries from April 1, 2023 to October 15, 2023**

| PANGO lineage | Relative $R_e$ (posterior value) | | | $\hat{R}$ | Bulk effective sample size | Tail effective sample size |
|---|---|---|---|---|---|---|
| | Mean | $2.5^{th}$ percentile | $97.5^{th}$ percentile | | | |
| BA.2.86 | 1.331 | 1.280 | 1.396 | 1.000 | 12500.997 | 9231.915 |
| HK.3 | 1.290 | 1.276 | 1.304 | 1.000 | 15666.432 | 11978.857 |
| FL.1.5.1 | 1.188 | 1.172 | 1.206 | 1.000 | 17742.666 | 11532.106 |
| EG.5.1.1 | 1.164 | 1.153 | 1.174 | 1.000 | 18492.765 | 11344.029 |
| EG.5.1 | 1.154 | 1.142 | 1.167 | 1.000 | 18710.667 | 12009.460 |
| FU.1 | 1.082 | 1.070 | 1.094 | 1.001 | 22406.928 | 11507.884 |
| XBB.1.16 | 1.069 | 1.060 | 1.078 | 1.000 | 21920.865 | 11583.298 |
| XBB.1.16.1 | 1.066 | 1.056 | 1.076 | 1.000 | 20782.767 | 10678.185 |
| XBB.2.3 | 1.066 | 1.057 | 1.075 | 1.001 | 21395.539 | 11433.803 |
| EG.1 | 1.046 | 1.037 | 1.056 | 1.000 | 19588.427 | 11151.843 |
| FL.4 | 1.037 | 1.030 | 1.044 | 1.000 | 18390.207 | 11351.786 |
| FL.2 | 1.032 | 1.022 | 1.043 | 1.000 | 20026.417 | 12271.067 |
| XBB.1.9.1 | 1.032 | 1.026 | 1.037 | 1.000 | 18231.883 | 11745.673 |
| XBB.1.9.2 | 1.031 | 1.021 | 1.041 | 1.000 | 22296.467 | 11687.795 |
| FL.5 | 1.027 | 1.017 | 1.037 | 1.000 | 19027.682 | 12710.259 |
| XBB.1.5.24 | 0.999 | 0.987 | 1.010 | 1.000 | 18259.622 | 13047.822 |

**Table S3. Estimated national relative $R_e$ and related epidemic dynamics modeling parameters of the SARS-CoV-2 lineages of interest circulated in 13 countries from April 1, 2023 to October 15, 2023**

| | | Relative $R_e$ (posterior value) | | | | | |
|---|---|---|---|---|---|---|---|
| PANGO lineage | Country | Mean | $2.5^{th}$ percentile | $97.5^{th}$ percentile | $\hat{R}$ | Bulk effective sample size | Tail effective sample size |
| BA.2.86 | Australia | 1.292 | 1.194 | 1.399 | 1.000 | 16992.179 | 9888.423 |
| HK.3 | Australia | 1.298 | 1.274 | 1.326 | 1.000 | 16914.570 | 11025.461 |
| XBB.1.16 | Australia | 1.061 | 1.056 | 1.068 | 1.000 | 9521.602 | 11821.095 |
| XBB.1.16.1 | Australia | 1.062 | 1.054 | 1.069 | 1.000 | 12018.740 | 12599.905 |
| XBB.1.5.24 | Australia | 0.991 | 0.968 | 1.011 | 1.000 | 21110.155 | 12246.698 |
| XBB.1.9.1 | Australia | 1.034 | 1.025 | 1.042 | 1.000 | 14852.946 | 12198.298 |
| XBB.1.9.2 | Australia | 1.038 | 1.030 | 1.047 | 1.000 | 14829.864 | 12400.851 |
| XBB.2.3 | Australia | 1.054 | 1.044 | 1.063 | 1.000 | 14723.119 | 13300.255 |
| EG.1 | Australia | 1.038 | 1.031 | 1.045 | 1.000 | 11801.002 | 12419.321 |
| EG.5.1 | Australia | 1.160 | 1.149 | 1.171 | 1.000 | 12877.574 | 11582.042 |
| EG.5.1.1 | Australia | 1.180 | 1.169 | 1.192 | 1.000 | 12749.206 | 11542.987 |
| FL.1.5.1 | Australia | 1.223 | 1.195 | 1.256 | 1.000 | 18241.736 | 11351.934 |
| FL.2 | Australia | 1.040 | 1.027 | 1.053 | 1.000 | 19550.899 | 12602.324 |
| FL.4 | Australia | 1.045 | 1.034 | 1.057 | 1.000 | 17246.029 | 12008.466 |
| FL.5 | Australia | 1.047 | 1.026 | 1.068 | 1.000 | 19011.405 | 13234.591 |
| FU.1 | Australia | 1.081 | 1.067 | 1.094 | 1.000 | 16746.135 | 12825.198 |
| BA.2.86 | Spain | 1.343 | 1.299 | 1.393 | 1.000 | 23899.233 | 11804.195 |
| HK.3 | Spain | 1.296 | 1.272 | 1.322 | 1.000 | 18999.891 | 11431.681 |
| XBB.1.16 | Spain | 1.080 | 1.075 | 1.086 | 1.000 | 14338.805 | 13285.254 |
| XBB.1.16.1 | Spain | 1.075 | 1.066 | 1.084 | 1.000 | 20462.708 | 12683.609 |
| XBB.1.5.24 | Spain | 0.985 | 0.961 | 1.005 | 1.000 | 19069.138 | 12363.094 |
| XBB.1.9.1 | Spain | 1.027 | 1.021 | 1.033 | 1.000 | 18731.971 | 12416.789 |
| XBB.1.9.2 | Spain | 1.014 | 1.007 | 1.021 | 1.000 | 21249.213 | 12958.060 |
| XBB.2.3 | Spain | 1.047 | 1.041 | 1.053 | 1.000 | 17823.176 | 13072.619 |
| EG.1 | Spain | 1.026 | 1.018 | 1.033 | 1.000 | 21266.527 | 12805.227 |
| EG.5.1 | Spain | 1.156 | 1.147 | 1.166 | 1.000 | 15205.154 | 12227.235 |
| EG.5.1.1 | Spain | 1.185 | 1.173 | 1.197 | 1.000 | 15968.266 | 11821.796 |
| FL.1.5.1 | Spain | 1.179 | 1.165 | 1.195 | 1.000 | 20382.618 | 11967.645 |
| FL.2 | Spain | 1.033 | 1.019 | 1.047 | 1.000 | 25950.882 | 11701.514 |
| FL.4 | Spain | 1.034 | 1.028 | 1.041 | 1.000 | 18578.221 | 12301.054 |
| FL.5 | Spain | 1.012 | 0.998 | 1.025 | 1.000 | 21131.773 | 12498.158 |
| FU.1 | Spain | 1.092 | 1.078 | 1.107 | 1.000 | 25034.969 | 12019.345 |
| BA.2.86 | Sweden | 1.268 | 1.225 | 1.313 | 1.000 | 19382.521 | 12573.476 |
| HK.3 | Sweden | 1.301 | 1.274 | 1.333 | 1.000 | 16348.086 | 10375.703 |
| XBB.1.16 | Sweden | 1.073 | 1.065 | 1.081 | 1.000 | 13252.305 | 13513.698 |
| XBB.1.16.1 | Sweden | 1.048 | 1.038 | 1.058 | 1.000 | 15929.427 | 12805.816 |
| XBB.1.5.24 | Sweden | 1.015 | 1.002 | 1.029 | 1.000 | 18687.337 | 12433.456 |
| XBB.1.9.1 | Sweden | 1.029 | 1.022 | 1.037 | 1.000 | 15825.425 | 12822.334 |
| XBB.1.9.2 | Sweden | 1.042 | 1.031 | 1.053 | 1.000 | 18421.575 | 12882.507 |
| XBB.2.3 | Sweden | 1.077 | 1.065 | 1.090 | 1.000 | 17300.610 | 11944.894 |
| EG.1 | Sweden | 1.036 | 1.026 | 1.046 | 1.001 | 18827.869 | 12801.802 |
| EG.5.1 | Sweden | 1.193 | 1.176 | 1.212 | 1.000 | 13054.777 | 12322.789 |
| EG.5.1.1 | Sweden | 1.182 | 1.169 | 1.196 | 1.000 | 11466.005 | 10765.362 |
| FL.1.5.1 | Sweden | 1.209 | 1.185 | 1.236 | 1.000 | 15327.037 | 10775.758 |

| | | | | | | | |
|---|---|---|---|---|---|---|---|
| FL.2 | Sweden | 1.032 | 1.015 | 1.048 | 1.001 | 22188.219 | 12875.997 |
| FL.4 | Sweden | 1.038 | 1.027 | 1.048 | 1.000 | 17500.550 | 12599.755 |
| FL.5 | Sweden | 1.043 | 1.027 | 1.059 | 1.000 | 21161.065 | 13141.231 |
| FU.1 | Sweden | 1.095 | 1.082 | 1.109 | 1.000 | 17186.916 | 12750.820 |
| BA.2.86 | United Kingdom | 1.257 | 1.235 | 1.280 | 1.001 | 18363.120 | 11310.604 |
| HK.3 | United Kingdom | 1.279 | 1.258 | 1.299 | 1.000 | 19080.570 | 12372.706 |
| XBB.1.16 | United Kingdom | 1.085 | 1.081 | 1.090 | 1.000 | 9244.688 | 11010.834 |
| XBB.1.16.1 | United Kingdom | 1.074 | 1.068 | 1.080 | 1.000 | 12323.686 | 11872.917 |
| XBB.1.5.24 | United Kingdom | 1.001 | 0.985 | 1.016 | 1.001 | 25317.014 | 13176.982 |
| XBB.1.9.1 | United Kingdom | 1.022 | 1.016 | 1.028 | 1.000 | 15054.861 | 11173.187 |
| XBB.1.9.2 | United Kingdom | 1.025 | 1.018 | 1.031 | 1.000 | 16573.473 | 13401.091 |
| XBB.2.3 | United Kingdom | 1.069 | 1.062 | 1.076 | 1.000 | 14414.014 | 12844.793 |
| EG.1 | United Kingdom | 1.049 | 1.042 | 1.056 | 1.000 | 15252.430 | 12619.099 |
| EG.5.1 | United Kingdom | 1.152 | 1.144 | 1.160 | 1.000 | 12071.703 | 12430.057 |
| EG.5.1.1 | United Kingdom | 1.150 | 1.143 | 1.158 | 1.000 | 11495.224 | 11348.895 |
| FL.1.5.1 | United Kingdom | 1.168 | 1.158 | 1.179 | 1.000 | 14355.727 | 11727.959 |
| FL.2 | United Kingdom | 1.031 | 1.020 | 1.042 | 1.000 | 21210.683 | 12286.419 |
| FL.4 | United Kingdom | 1.055 | 1.047 | 1.062 | 1.001 | 15397.208 | 12713.877 |
| FL.5 | United Kingdom | 1.025 | 1.014 | 1.036 | 1.000 | 22873.037 | 13273.923 |
| FU.1 | United Kingdom | 1.093 | 1.083 | 1.103 | 1.000 | 17827.346 | 12920.765 |
| BA.2.86 | USA | 1.263 | 1.230 | 1.299 | 1.000 | 25262.836 | 11297.788 |
| HK.3 | USA | 1.282 | 1.269 | 1.295 | 1.000 | 24623.411 | 12489.115 |
| XBB.1.16 | USA | 1.068 | 1.067 | 1.070 | 1.000 | 13585.547 | 12089.200 |
| XBB.1.16.1 | USA | 1.067 | 1.065 | 1.069 | 1.000 | 16178.103 | 12211.922 |
| XBB.1.5.24 | USA | 1.011 | 1.003 | 1.020 | 1.000 | 25061.372 | 11985.283 |
| XBB.1.9.1 | USA | 1.040 | 1.038 | 1.043 | 1.000 | 21820.863 | 13318.741 |
| XBB.1.9.2 | USA | 1.041 | 1.038 | 1.045 | 1.000 | 21777.164 | 12537.105 |
| XBB.2.3 | USA | 1.079 | 1.076 | 1.081 | 1.000 | 16176.639 | 12696.166 |
| EG.1 | USA | 1.049 | 1.046 | 1.053 | 1.000 | 22684.900 | 12672.595 |
| EG.5.1 | USA | 1.131 | 1.128 | 1.134 | 1.000 | 14470.705 | 12685.299 |
| EG.5.1.1 | USA | 1.148 | 1.144 | 1.151 | 1.000 | 14953.895 | 11997.165 |
| FL.1.5.1 | USA | 1.166 | 1.162 | 1.170 | 1.000 | 14389.195 | 11073.154 |
| FL.2 | USA | 1.037 | 1.032 | 1.041 | 1.000 | 25616.530 | 13342.207 |
| FL.4 | USA | 1.044 | 1.041 | 1.048 | 1.000 | 24359.508 | 11747.569 |
| FL.5 | USA | 1.039 | 1.033 | 1.044 | 1.000 | 24282.527 | 13246.614 |
| FU.1 | USA | 1.075 | 1.072 | 1.078 | 1.000 | 19468.036 | 13055.833 |
| BA.2.86 | Austria | 1.370 | 1.219 | 1.639 | 1.000 | 10107.914 | 5341.746 |
| HK.3 | Austria | 1.260 | 1.213 | 1.298 | 1.000 | 13209.036 | 12240.729 |
| XBB.1.16 | Austria | 1.085 | 1.075 | 1.096 | 1.000 | 18375.829 | 12729.044 |
| XBB.1.16.1 | Austria | 1.081 | 1.066 | 1.098 | 1.000 | 20811.977 | 13082.452 |
| XBB.1.5.24 | Austria | 0.992 | 0.968 | 1.013 | 1.000 | 22295.578 | 12937.962 |
| XBB.1.9.1 | Austria | 1.037 | 1.028 | 1.047 | 1.000 | 22291.966 | 12948.751 |
| XBB.1.9.2 | Austria | 1.020 | 1.004 | 1.035 | 1.000 | 21514.641 | 12613.120 |
| XBB.2.3 | Austria | 1.078 | 1.064 | 1.092 | 1.000 | 22234.678 | 12930.704 |
| EG.1 | Austria | 1.053 | 1.042 | 1.065 | 1.000 | 20986.223 | 13268.598 |
| EG.5.1 | Austria | 1.159 | 1.140 | 1.179 | 1.000 | 20736.896 | 11672.283 |
| EG.5.1.1 | Austria | 1.169 | 1.154 | 1.185 | 1.000 | 18661.981 | 12323.562 |
| FL.1.5.1 | Austria | 1.192 | 1.168 | 1.221 | 1.000 | 18074.480 | 10261.511 |
| FL.2 | Austria | 1.032 | 1.015 | 1.049 | 1.000 | 25472.454 | 13112.791 |

| | | | | | | | |
|---|---|---|---|---|---|---|---|
| FL.4 | Austria | 1.025 | 1.005 | 1.040 | 1.000 | 19515.867 | 11898.808 |
| FL.5 | Austria | 1.027 | 1.007 | 1.048 | 1.000 | 24684.505 | 12292.823 |
| FU.1 | Austria | 1.086 | 1.069 | 1.103 | 1.000 | 20831.031 | 12842.205 |
| BA.2.86 | Canada | 1.354 | 1.299 | 1.416 | 1.000 | 20403.045 | 11167.707 |
| HK.3 | Canada | 1.300 | 1.283 | 1.318 | 1.001 | 19426.054 | 12168.027 |
| XBB.1.16 | Canada | 1.062 | 1.059 | 1.066 | 1.000 | 11078.891 | 12535.608 |
| XBB.1.16.1 | Canada | 1.066 | 1.062 | 1.071 | 1.000 | 15422.020 | 12544.744 |
| XBB.1.5.24 | Canada | 1.001 | 0.982 | 1.018 | 1.000 | 23389.733 | 12076.777 |
| XBB.1.9.1 | Canada | 1.035 | 1.030 | 1.040 | 1.000 | 18439.982 | 12994.520 |
| XBB.1.9.2 | Canada | 1.042 | 1.034 | 1.050 | 1.000 | 19391.923 | 10892.606 |
| XBB.2.3 | Canada | 1.064 | 1.059 | 1.070 | 1.000 | 15761.384 | 12419.310 |
| EG.1 | Canada | 1.048 | 1.041 | 1.055 | 1.000 | 18866.448 | 12037.743 |
| EG.5.1 | Canada | 1.151 | 1.144 | 1.157 | 1.000 | 12291.486 | 11115.280 |
| EG.5.1.1 | Canada | 1.147 | 1.141 | 1.152 | 1.000 | 10881.593 | 12453.035 |
| FL.1.5.1 | Canada | 1.177 | 1.168 | 1.185 | 1.001 | 14346.830 | 12606.126 |
| FL.2 | Canada | 1.060 | 1.050 | 1.071 | 1.000 | 20171.440 | 12475.257 |
| FL.4 | Canada | 1.035 | 1.028 | 1.042 | 1.000 | 23877.028 | 11677.903 |
| FL.5 | Canada | 1.024 | 1.015 | 1.033 | 1.000 | 22268.850 | 12472.768 |
| FU.1 | Canada | 1.107 | 1.099 | 1.115 | 1.000 | 17939.871 | 11845.472 |
| BA.2.86 | China | 1.345 | 1.232 | 1.497 | 1.000 | 19929.112 | 10179.424 |
| HK.3 | China | 1.315 | 1.303 | 1.327 | 1.000 | 7556.564 | 10566.592 |
| XBB.1.16 | China | 1.062 | 1.055 | 1.069 | 1.000 | 5759.088 | 8538.200 |
| XBB.1.16.1 | China | 1.077 | 1.068 | 1.086 | 1.000 | 7681.159 | 10560.075 |
| XBB.1.5.24 | China | 1.014 | 0.997 | 1.031 | 1.000 | 18012.468 | 10538.449 |
| XBB.1.9.1 | China | 1.038 | 1.030 | 1.046 | 1.000 | 8759.527 | 10140.351 |
| XBB.1.9.2 | China | 1.043 | 1.029 | 1.058 | 1.000 | 16389.151 | 12258.435 |
| XBB.2.3 | China | 1.067 | 1.051 | 1.083 | 1.000 | 18056.657 | 12536.969 |
| EG.1 | China | 1.076 | 1.057 | 1.094 | 1.000 | 15900.882 | 12204.677 |
| EG.5.1 | China | 1.163 | 1.153 | 1.173 | 1.001 | 7450.557 | 9776.909 |
| EG.5.1.1 | China | 1.164 | 1.157 | 1.171 | 1.001 | 4860.004 | 7617.604 |
| FL.1.5.1 | China | 1.199 | 1.173 | 1.228 | 1.000 | 18422.742 | 11089.515 |
| FL.2 | China | 1.029 | 1.020 | 1.037 | 1.000 | 9476.306 | 11742.658 |
| FL.4 | China | 1.048 | 1.041 | 1.055 | 1.001 | 6657.045 | 9228.329 |
| FL.5 | China | 1.019 | 1.000 | 1.036 | 1.000 | 19353.932 | 11122.307 |
| FU.1 | China | 1.070 | 1.064 | 1.077 | 1.001 | 5159.778 | 8699.827 |
| BA.2.86 | France | 1.393 | 1.347 | 1.444 | 1.000 | 18057.777 | 10613.750 |
| HK.3 | France | 1.298 | 1.276 | 1.323 | 1.000 | 16488.167 | 11010.176 |
| XBB.1.16 | France | 1.086 | 1.080 | 1.092 | 1.000 | 11178.216 | 12370.373 |
| XBB.1.16.1 | France | 1.071 | 1.064 | 1.078 | 1.000 | 15214.628 | 12868.528 |
| XBB.1.5.24 | France | 0.996 | 0.978 | 1.011 | 1.000 | 24108.228 | 12796.105 |
| XBB.1.9.1 | France | 1.031 | 1.026 | 1.037 | 1.000 | 15180.194 | 12845.606 |
| XBB.1.9.2 | France | 1.021 | 1.014 | 1.029 | 1.001 | 21182.179 | 13293.139 |
| XBB.2.3 | France | 1.081 | 1.073 | 1.089 | 1.000 | 15365.387 | 12421.625 |
| EG.1 | France | 1.038 | 1.032 | 1.045 | 1.000 | 15356.731 | 12510.441 |
| EG.5.1 | France | 1.171 | 1.161 | 1.181 | 1.000 | 11960.115 | 10953.079 |
| EG.5.1.1 | France | 1.163 | 1.155 | 1.172 | 1.000 | 11115.199 | 12287.407 |
| FL.1.5.1 | France | 1.192 | 1.180 | 1.206 | 1.000 | 13728.181 | 11489.558 |
| FL.2 | France | 1.021 | 1.009 | 1.032 | 1.000 | 22068.045 | 12509.221 |
| FL.4 | France | 1.028 | 1.021 | 1.036 | 1.000 | 19169.265 | 12711.607 |
| FL.5 | France | 1.012 | 1.001 | 1.022 | 1.000 | 21408.237 | 13313.552 |

| | | | | | | | |
|---|---|---|---|---|---|---|---|
| FU.1 | France | 1.097 | 1.085 | 1.109 | 1.000 | 20276.473 | 12473.355 |
| BA.2.86 | Italy | 1.384 | 1.249 | 1.613 | 1.000 | 11121.400 | 7404.732 |
| HK.3 | Italy | 1.291 | 1.260 | 1.326 | 1.001 | 17247.156 | 10905.489 |
| XBB.1.16 | Italy | 1.072 | 1.064 | 1.081 | 1.000 | 15174.779 | 12710.843 |
| XBB.1.16.1 | Italy | 1.085 | 1.070 | 1.101 | 1.000 | 18400.405 | 11551.858 |
| XBB.1.5.24 | Italy | 0.997 | 0.972 | 1.018 | 1.000 | 21480.698 | 12394.726 |
| XBB.1.9.1 | Italy | 1.027 | 1.019 | 1.034 | 1.001 | 15769.599 | 12721.921 |
| XBB.1.9.2 | Italy | 1.021 | 1.009 | 1.032 | 1.000 | 20612.665 | 12264.219 |
| XBB.2.3 | Italy | 1.075 | 1.066 | 1.085 | 1.000 | 15240.025 | 12445.643 |
| EG.1 | Italy | 1.025 | 1.016 | 1.034 | 1.000 | 16808.054 | 12144.522 |
| EG.5.1 | Italy | 1.149 | 1.136 | 1.163 | 1.000 | 16435.776 | 11580.215 |
| EG.5.1.1 | Italy | 1.164 | 1.150 | 1.178 | 1.000 | 15886.685 | 11639.311 |
| FL.1.5.1 | Italy | 1.165 | 1.143 | 1.187 | 1.000 | 20498.120 | 12607.563 |
| FL.2 | Italy | 1.028 | 1.012 | 1.043 | 1.000 | 21726.978 | 10864.276 |
| FL.4 | Italy | 1.025 | 1.008 | 1.039 | 1.000 | 18738.713 | 11328.998 |
| FL.5 | Italy | 1.034 | 1.021 | 1.047 | 1.000 | 20579.173 | 13437.995 |
| FU.1 | Italy | 1.062 | 1.045 | 1.078 | 1.000 | 21658.501 | 12592.094 |
| BA.2.86 | Japan | 1.335 | 1.278 | 1.400 | 1.000 | 21857.443 | 11349.725 |
| HK.3 | Japan | 1.265 | 1.246 | 1.284 | 1.000 | 17074.136 | 12295.305 |
| XBB.1.16 | Japan | 1.041 | 1.038 | 1.045 | 1.001 | 7569.017 | 10705.486 |
| XBB.1.16.1 | Japan | 1.030 | 1.026 | 1.035 | 1.001 | 10102.233 | 11813.774 |
| XBB.1.5.24 | Japan | 0.980 | 0.968 | 0.991 | 1.000 | 21750.292 | 13442.419 |
| XBB.1.9.1 | Japan | 1.022 | 1.017 | 1.026 | 1.000 | 11305.319 | 11972.928 |
| XBB.1.9.2 | Japan | 1.045 | 1.039 | 1.051 | 1.000 | 15049.714 | 12019.519 |
| XBB.2.3 | Japan | 1.047 | 1.041 | 1.053 | 1.000 | 13021.250 | 11814.756 |
| EG.1 | Japan | 1.048 | 1.043 | 1.053 | 1.001 | 11323.399 | 11890.000 |
| EG.5.1 | Japan | 1.119 | 1.115 | 1.124 | 1.000 | 8919.943 | 11396.780 |
| EG.5.1.1 | Japan | 1.135 | 1.130 | 1.140 | 1.000 | 9289.792 | 10874.983 |
| FL.1.5.1 | Japan | 1.169 | 1.142 | 1.196 | 1.000 | 20360.026 | 11888.688 |
| FL.2 | Japan | 1.013 | 1.007 | 1.019 | 1.000 | 15427.076 | 12697.171 |
| FL.4 | Japan | 1.032 | 1.027 | 1.037 | 1.000 | 11123.278 | 12072.479 |
| FL.5 | Japan | 1.013 | 1.005 | 1.021 | 1.000 | 17295.122 | 11555.509 |
| FU.1 | Japan | 1.049 | 1.043 | 1.055 | 1.000 | 13543.550 | 11987.560 |
| BA.2.86 | Singapore | 1.456 | 1.289 | 1.836 | 1.000 | 7694.237 | 6085.526 |
| HK.3 | Singapore | 1.293 | 1.277 | 1.311 | 1.000 | 11252.488 | 11122.279 |
| XBB.1.16 | Singapore | 1.051 | 1.042 | 1.061 | 1.000 | 9646.400 | 12008.663 |
| XBB.1.16.1 | Singapore | 1.052 | 1.043 | 1.062 | 1.000 | 9010.364 | 11777.741 |
| XBB.1.5.24 | Singapore | 0.990 | 0.965 | 1.011 | 1.000 | 17812.931 | 11803.787 |
| XBB.1.9.1 | Singapore | 1.026 | 1.013 | 1.038 | 1.000 | 17432.875 | 11652.534 |
| XBB.1.9.2 | Singapore | 1.000 | 0.983 | 1.016 | 1.000 | 15539.160 | 13005.707 |
| XBB.2.3 | Singapore | 1.050 | 1.035 | 1.065 | 1.000 | 14731.766 | 12359.817 |
| EG.1 | Singapore | 1.053 | 1.034 | 1.075 | 1.000 | 21878.278 | 11587.127 |
| EG.5.1 | Singapore | 1.166 | 1.150 | 1.184 | 1.000 | 13285.534 | 11607.936 |
| EG.5.1.1 | Singapore | 1.170 | 1.159 | 1.182 | 1.000 | 9035.052 | 10125.679 |
| FL.1.5.1 | Singapore | 1.225 | 1.196 | 1.259 | 1.000 | 16002.130 | 11628.369 |
| FL.2 | Singapore | 1.018 | 0.995 | 1.037 | 1.000 | 20009.470 | 12551.398 |
| FL.4 | Singapore | 1.037 | 1.025 | 1.049 | 1.000 | 15293.533 | 11579.466 |
| FL.5 | Singapore | 1.019 | 0.999 | 1.036 | 1.001 | 19626.726 | 11924.557 |
| FU.1 | Singapore | 1.059 | 1.048 | 1.069 | 1.000 | 10138.594 | 12756.287 |
| BA.2.86 | South Korea | 1.328 | 1.246 | 1.430 | 1.000 | 20160.244 | 10103.431 |

| | | | | | | | |
|---|---|---|---|---|---|---|---|
| HK.3 | South Korea | 1.284 | 1.269 | 1.298 | 1.000 | 17096.312 | 12563.696 |
| XBB.1.16 | South Korea | 1.068 | 1.063 | 1.072 | 1.000 | 7320.100 | 10412.781 |
| XBB.1.16.1 | South Korea | 1.056 | 1.050 | 1.063 | 1.001 | 12513.196 | 12238.227 |
| XBB.1.5.24 | South Korea | 1.017 | 1.006 | 1.029 | 1.000 | 21830.588 | 11788.863 |
| XBB.1.9.1 | South Korea | 1.044 | 1.039 | 1.050 | 1.000 | 9791.179 | 12370.272 |
| XBB.1.9.2 | South Korea | 1.038 | 1.032 | 1.043 | 1.000 | 10593.534 | 11121.287 |
| XBB.2.3 | South Korea | 1.065 | 1.059 | 1.071 | 1.000 | 10830.279 | 11212.136 |
| EG.1 | South Korea | 1.066 | 1.061 | 1.070 | 1.000 | 8139.919 | 10111.124 |
| EG.5.1 | South Korea | 1.138 | 1.133 | 1.144 | 1.001 | 7541.307 | 10568.760 |
| EG.5.1.1 | South Korea | 1.167 | 1.161 | 1.173 | 1.000 | 8227.120 | 11485.626 |
| FL.1.5.1 | South Korea | 1.195 | 1.166 | 1.229 | 1.000 | 20144.319 | 10584.435 |
| FL.2 | South Korea | 1.055 | 1.047 | 1.062 | 1.000 | 12967.409 | 10665.172 |
| FL.4 | South Korea | 1.038 | 1.031 | 1.046 | 1.000 | 14742.428 | 12272.465 |
| FL.5 | South Korea | 1.035 | 1.025 | 1.046 | 1.000 | 16694.108 | 12240.837 |
| FU.1 | South Korea | 1.086 | 1.079 | 1.094 | 1.000 | 12701.385 | 10343.798 |

**Table S4. Primers used in this study**

| Primer name | Primer sequence (5'-to-3') | Purpose |
|---|---|---|
| Omicron universal Fw | cactatagggcgaattgggtaccatgtttgtgttcctggt | Preparation of S expression plasmid |
| BA.2 WT Rv | agctccaccgcggtggcggccgctcaggtgtagtgcagtttca | Preparation of S expression plasmid |
| pC-S-XBB.1.5 _F445L_Fwd | caactacctctacagaTTCttcaggaagagcaagctg | Preparation of S expression plasmid |
| pC-S-XBB.1.5 _F455L_Rev | cagcttgctcttcctgaaGAAtctgtagaggtagttg | Preparation of S expression plasmid |
| pC-S-XBB.1.5 _FL445-456LF_Fwd | caactacctctacagaTTCCTGaggaagagcaagctg | Preparation of S expression plasmid |
| pC-S-XBB.1.5 _FL445-456LF_Rev | cagcttgctcttcctCAGGAAtctgtagaggtagttg | Preparation of S expression plasmid |

### Consortia

#### The Genotype to Phenotype Japan (G2P-Japan) Consortium

**The Institute of Medical Science, The University of Tokyo, Japan**
Naoko Misawa, Ziyi Guo, Jarel Elgin M. Tolentino, Shigeru Fujita, Lin Pan, Mai Suganami, Mika Chiba, Ryo Yoshimura, Kyoko Yasuda, Keiko Iida, Naomi Ohsumi, Adam P. Strange, Shiho Tanaka

**Hokkaido University, Japan**
Takasuke Fukuhara, Tomokazu Tamura, Rigel Suzuki, Saori Suzuki, Hayato Ito, Keita Matsuno, Hirofumi Sawa, Naganori Nao, Shinya Tanaka, Masumi Tsuda, Lei Wang, Yoshikata Oda, Zannatul Ferdous, Kenji Shishido

**Tokai University, Japan**
So Nakagawa

**Kyoto University, Japan**
Kotaro Shirakawa, Akifumi Takaori-Kondo, Kayoko Nagata, Ryosuke Nomura, Yoshihito Horisawa, Yusuke Tashiro, Yugo Kawai, Kazuo Takayama, Rina Hashimoto, Sayaka Deguchi, Yukio Watanabe, Ayaka Sakamoto, Naoko Yasuhara, Takao Hashiguchi, Tateki Suzuki, Kanako Kimura, Jiei Sasaki, Yukari Nakajima, Hisano Yajima

**Hiroshima University, Japan**
Takashi Irie, Ryoko Kawabata

**Kyushu University, Japan**
Kaori Tabata

**Kumamoto University, Japan**
Terumasa Ikeda, Hesham Nasser, Ryo Shimizu, MST Monira Begum, Michael Jonathan, Yuka Mugita, Otowa Takahashi, Kimiko Ichihara, Takamasa Ueno, Chihiro Motozono, Mako Toyoda

**University of Miyazaki, Japan**
Akatsuki Saito, Maya Shofa, Yuki Shibatani, Tomoko Nishiuchi

**Acknowledgments**

We would like to thank all members of The Genotype to Phenotype Japan (G2P-Japan) Consortium. We thank Dr. Kenzo Tokunaga (National Institute of Infectious Diseases, Japan) for sharing materials. We gratefully acknowledge the numerous laboratories worldwide that have provided sequence data and metadata to GISAID. A full list of originating and submitting laboratories for the sequences used in our analysis can be found at https://www.gisaid.org using the EPI-SET-ID: EPI_SET_231110rt and EPI_SET_231111fo.
